# BroodScan: A Honeybee Brood Scanner Technology

**DOI:** 10.1101/2025.05.17.651227

**Authors:** Martin Stefanec, Daniel N. Hofstadler, Martin Kärcher, Thomas Schmickl

## Abstract

Observing honeybee (*Apis mellifera*) brood dynamics and host-parasite interactions, especially within sealed cells, is vital for colony health research but traditionally relies on disruptive or destructive methods. Here we present BroodScan, a monitoring system employing a modified consumer-grade Charge-Coupled Device (CCD) flatbed scanner for continuous, high-resolution basal imaging of brood cells directly within an active beehive. Our experiments, conducted over a 35-day period, demonstrate BroodScan’s capacity to track the complete honeybee brood development cycle from egg to adult emergence, yielding quantitative data such as an average egg stage duration of 72.93 ± 2.04 hours (n=58). Notably, the system provided detailed, time-lapse visual documentation of the complete intra-cellular reproductive cycle of the parasitic mite *Varroa destructor* within sealed worker cells, including the progression from foundress mite to multiple distinct offspring stages and their maturation. BroodScan offers a valuable, non-invasive tool that overcomes key limitations of previous techniques, opening new avenues for research into honeybee biology, colony health, and the impacts of diverse stressors like *Varroa destructor*, thereby promising to enhance our understanding and stewardship of these vital pollinators.

## 1 INTRODUCTION

### 1.1 Motivation

Honeybees hold immense agricultural, ecological and socio-economic significance, both through their direct products (honey, pollen, propolis & wax) with honey alone valued globally at approximately USD 9.41 billion in 2023 [1]. Their indirect service of pollination is also paramount; while the total annual global food production reliant on animal pollinators (including bees, native bees, and other insects) is valued around EUR 625 billion [2], honeybees are recognized as key contributors to this essential service [3]. This impact is largely attributed to the eusocial organization of honeybee colonies, where individual behaviors culminate in a highly complex, self-organized system, facilitating resource allocation, energy balance, regulation of the hive environment, and swift responses to environmental threats. Despite this, colonies face unprecedented challenges, with 20-50% [4–7] experiencing annual losses due to various diseases and stressors, many of which affect the growth, survival, or health of the brood.

Our motivation to develop new technology to better study the brood nest of a honeybee colony stems from two interconnected aspects: (1) the significant impact of honeybee stressors within the brood nest area, where the majority of their detrimental factors unfold [8–10], and (2) the pivotal role of this region as the sole origin of 100% of the colony’s population, including the workers vital for pollination services and colony products [11].

### 1.2 Challenges

Unfortunately, the brood nest’s dynamics, economics, and morphogenesis are significantly under-researched due to the intricate networks of self-regulatory mechanisms involved in nurturing the brood, preparing cells, egg laying, and managing nutrient logistics. Understanding this complexity will provide essential insights into breeding, managing, and protecting honeybee colonies.

In consequence, the brood nest presents a challenging yet compelling subject for study due to its emergent complexity, characterized by cascading feedback loops of self-regulation and self-organization. Our technological observation approach aims at generating a unique “big data” dataset in the long-term, most favorable across a whole honeybee brood season (spring to autumn). With such technology, newfound insights will have the potential to better protect honeybee colonies by identifying the impacts and vulnerabilities of their stressors, thereby serving as a foundation for safeguarding these crucial pollinators.

These observational challenges are further complicated by the limitations of traditional methods for studying these developmental processes. Direct observation often requires the removal of brood frames from the hive, a process that can disrupt the thermal regulation and social organization of the colony, potentially altering the natural developmental or behavioral processes being studied [12–14]. These challenges underscore the need for less invasive and more continuous monitoring techniques. The adaptation of existing technologies, particularly from outside traditional biology, offers one avenue for innovation. For instance, flatbed scanner technology, originally designed for document digitization, has been increasingly repurposed for diverse biological applications due to its high resolution and affordability [15]. However, its application to dynamic, in-hive environments has specific hurdles. Furthermore, even direct observation of the comb surface is limited. Once a larva has completed its feeding and growth phase and begins to prepare for pupation, worker bees seal the brood cell with a wax cap. From this point, the developing bee inside is inaccessible to direct visual observation from above without resorting to destructive techniques such as manually uncapping cells or cutting the comb [13, 16]. This inability to see inside capped cells is a significant limitation, especially when trying to observe processes that occur specifically during this hidden period. For example, the reproductive cycle of the parasitic mite *Varroa destructor*, a major threat to honeybee health, takes place primarily within the sealed brood cell, making it extremely difficult to study in situ.

### 1.3 Honeybee Brood Biology

Many insects, including honeybees (*Apis mellifera* L.), undergo holometabolous development - a complete metamorphosis involving four distinct life stages: egg, larva, pupa and adult, with larval growth occurring over several instars after hatching [14]. Although it represents this common insect developmental pathway, honeybee larval development is unique because of its remarkable speed and consistency compared to that of many other insect species [17]. This rapid and highly predictable progression from egg to adult takes place within the precisely thermoregulated environment of the hive (maintained at approximately 33-36 °C [18], which largely insulates development from the direct influence of external climatic variability that governs developmental rates in many solitary insects.

The physiological development of honeybees is linked to their eusocial self-organization and self-regulation, which unfolds within a unique system of collective brood care [19]. Sister-nurse bees collectively care for the larvae, maintain cell hygiene and regulate local conditions, in complete contrast to the individual parental care strategies seen elsewhere in the animal kingdom [20]. Studying the inner workings of this rapid, consistent and socially managed individual development process provides insights into the biology of eusocial insects, colony resilience and responses to environmental cues. However, the very nature of the honeybee colony presents a challenge to *in situ* observations.

The brood nest is typically located at the center of the hive, physically protected and thermally insulated by the surrounding honeycombs and the colony [21]. After hatching, larvae develop within the same cells, dependent on nurse bee care [22]. However, observational studies by direct observation through the cell openings are often problematic: Even in observation hives, which are tailor made for observing the bees, the surface of the brood comb is often covered by a dense, mobile population of worker bees engaged in brood care, cell inspection or traversing the comb, preventing a continuous view of events in the cells below by blocking the line of sight. A typical frontal view of a honeycomb is shown in Fig. 1C. Adult bees often form layers on the comb surface, obscuring cell openings, while wax cappings further conceal the contents of brood or honey cells. The spatio-temporal dynamics of cell occupancy within a honeycomb are inherently complex due to the diverse and changing contents, ranging from brood at various developmental stages (egg, larva, pupa) to different forms of stored food (pollen, nectar, uncapped and capped honey), as shown in Fig. 1D.

**Figure 1:**
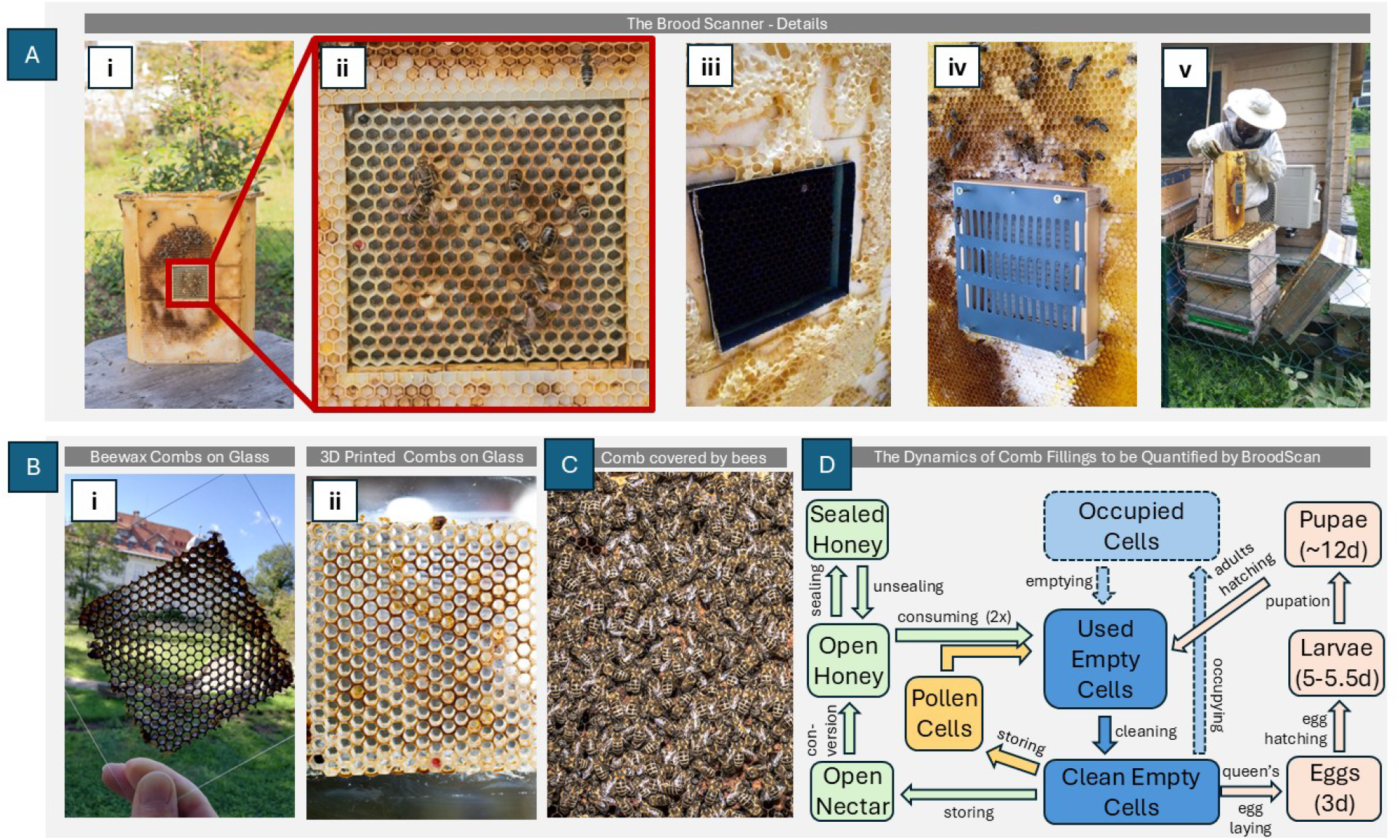
Overview of the BroodScan (A) Photos of the device and its parts. (i) Full device. (ii) Zoomed view of the 3D printed cells used in the experiments described here. (iii) The scan window without the 3D printed cells. (iv) A queen confinement cage to keep the queen for initial egg laying in the scanning window area. (v) Application of the BroodScan in a conventional box hive (Zander dimensions). (B) Applying combs on the scanner windows. (i) Example for bee-produced wax comb cells. (ii) Example of 3D printed comb cells. (C) Typical frontal view of a honeycomb. Direct visual access to internal cell contents is obstructed by adult bees covering the comb surface, often in multiple layers, and by the wax cappings that seal brood and honey cells. (D) Schematic representation of the dynamics of cell state transitions that BroodScan aims to observe and quantify.

### 1.4 The BroodScan Approach and Aims of this Paper

To address the challenges of observing intra-cellular dynamics within the honeybee brood nest, the BroodScan system was developed as part of the EU-funded Hiveopolis project (Grant Agreement No. 824069). The core technological concepts, involving a line scanner for in-hive basal observation of honeybee brood, were first formally documented in project deliverables starting in October 2022 [23]. Subsequent project reports further illustrated the system’s progression, including visual documentation of the technology and its establishment as a core project development (e.g., [24]; see also [25–27] for full developmental timeline).

The BroodScan technology, presented in this paper based on experiments conducted in 2022, critically employs a Charge-Coupled Device (CCD) based flatbed scanner and a distinct artificial comb interface. This approach prioritizes depth of field for high-resolution visualization of three-dimensional intra-cellular details. In this paper, we report on the detailed development and validation of the BroodScan system. We describe its key design innovations aimed at addressing major observational challenges in apiculture and present initial field data demonstrating its capability to track the complete honeybee brood cycle and visualize intra-cellular parasite dynamics.

### 1.5 Key Design Goals of BroodScan

With our technology, we have several design challenges that we faced and had to solve:

#### (1) Line of Sight

Honeybee comb observations, regardless of manual or automated, are usually performed in a top-down perspective using an observation hive. This means that cell contents are observed through the openings of the wax cells. This classic approach poses several severe problems: (1) Before pupation of a larva, the brood cell opening is closed by adult bees with a wax cap, thus there is no possible line of sight from above available anymore. (2) Even before this point, bees frequently crawl on the comb surface, or even crawl into the remaining empty cell space to feed or inspect the larva, this way also blocking the line of sight. (3) Not only brood cells, but also honey cells are often closed by the bees with a wax cap for long-term storage of the honey, impairing any top-down inspection system to visually inspect the stored honey. In order to avoid these problems, our BroodScan system is designed to observe the cell contents from the bottom side of the cells by applying specifically fabricated bottom-less comb cells (wax or 3D-printed) on a transparent base plate (glass or plexiglass), as seen in Fig. 1A-i-v. This way the line of sight can never be blocked by any cell caps or by any bees.

#### (2) Close-Distance Observations

The perspective from the bottom of the comb cell inherently provides the closest view for photographing brood development and nutrient accumulation (pollen, nectar, honey), which start at the cell base. Additionally, the close distance to the cell contents allows observation of details, such as body features or parasites.

#### (3) High-Resolution Imaging

For monitoring the growth and health of the brood, it is crucial to have high-resolution imaging across all observed cells, ideally across the whole brood comb. A flatbed scanner that can provide a high dpi resolution is an ideal solution for this requirement.

#### (4) Distortion-Free Observation

Another crucial issue with conventional top-down observation of comb cell contents is parallax: Depending on the camera lens’s focal length, only a limited number of cells may be visible down to their base. While parallax could be mitigated using longer focal length (telephoto) lenses, these require a greater distance of the camera from the comb, which is prevented by the tight space constraints within a standard hive. In addition to parallax, lens distortion also occurs with camera-based observations, typically correlated with the short focal length (wide-angle) lenses often needed for close work. A flatbed scanner avoids these issues, as this technology inherently produces images mostly free from both distortion and parallax.

#### (5) Applicability

The technology must have wide application potential, encompassing both physical integration and cost-effectiveness. Firstly, regarding the form factor, the device should ideally be installable in observation hives (which may require slight thickening to accommodate the unit while remaining functional) as well as conventional box hives, where space can be created by removing one or two adjacent frames. Secondly, regarding cost, the chosen approach must be affordable. Here too, a thin A4 or US-Letter flatbed scanner proves suitable, as its dimensions approximate a brood nest area, it is inexpensive, and it has a low profile.

#### (6) Long-Term Autonomy

For extended, minimally disruptive field deployment, the system is designed with low power consumption as a key goal. Ideally, the technology should support autonomous operation over long periods, potentially utilizing portable power sources like power banks or incorporating renewable energy harvesting (e.g., solar power) to minimize reliance on mains electricity.

#### (7) Simplicity

A key design goal was to create a device that is as “plug-and-play” as possible. Therefore, the primary approach utilizes 3D-printed comb sections mounted on a protective glass sheet placed on the scanner’s glass (see Fig. 1B-ii). This is considered the simplest, most versatile, and reusable method for providing the necessary comb structure for the brood nest. Simplicity also extended to the required software and electronics: a suitable flatbed scanner connects via USB to a single-board computer (SBC). The SBC operates on standard low-voltage (USB) power, stores images locally (e.g., on an SD card), and can transmit them via WiFi. Software control of the hardware is achieved via the scanner’s standard TWAIN interface [28], allowing simple scripts – for example, triggered by a “cron job” – to initiate scans.

#### (8) Minimal Disturbance

The measuring device must minimize interference with honeybee biology. Therefore, potential disturbances from scanning movement, excess heat, and illumination were considered. Scanning frequency should be minimized to limit disruption from movement. Excess heat from the scanner or its power supply may not be a major concern, as honeybees actively thermoregulate the brood nest to approximately 36–38 °C. Regarding illumination, future iterations could utilize red light, which is invisible to honeybees, to further reduce potential impact. However, preliminary experiments using white light, reported herein, showed no obvious detrimental effects on brood development. This lack of observed effect may potentially be explained by the developmental timeline of the honeybee visual system; photoreceptor and rhabdom maturation occurs throughout pupation, accelerating in the second half, but opsin expression levels increase dramatically only after adult emergence [29].

## 2 THE METHODOLOGY OF BROODSCAN

### 2.1 System Hardware and Design

The core imaging unit was an Epson Perfection V370 Photo flatbed scanner. This specific model was selected primarily because it utilizes Charge-Coupled Device (CCD) sensor technology [30] within a relatively compact A4-adjacent form factor; CCD provides essential depth of field for resolving details within honeycomb cells, a feature lacking in more common, thinner Contact Image Sensor (CIS) scanners [15]. While offering superior depth of field, CCD technology typically requires higher illumination levels than CIS, representing a potential trade-off concerning minimizing disturbance to the bees. Furthermore, while flatbed scanners significantly reduce the parallax distortion inherent in camera-based observations of honeycomb cells, preliminary tests indicated that this specific model still exhibited some parallax effects towards the lateral edges of its scanning area. Therefore, restricting image acquisition to the centered ROI (as described in Sec. 2.4, Fig 1A) was important for utilizing the area with the least parallax distortion, while also minimizing scan time, data volume, and potential light disturbance to the colony. The scanner operates using a dedicated 13.5 V DC power supply via its external AC adapter. Control and data transfer were managed by a single-board computer (SBC) (Raspberry Pi 4 model B) connected to the scanner via USB. The SBC ran Raspberry Pi OS (based on Debian 11 ‘Bullseye’) and the entire setup was installed within a Zander-type beehive.

To enable the scanner’s placement within a standard brood frame such that its surface could achieve direct contact with the comb, modifications were required to operate the scanner without its physical lid attached while maintaining the necessary electronic connection for initialization. The scanner base was housed within a custom-built wooden box (approx. 42 cm deep × 50 cm high × 7 cm wide, Fig. 1A-i,v), which was integrated into the hive structure. This box featured an aperture aligned with the scanning ROI located on the scanner’s platen; the other surfaces facing the colony were lined with plastic sheets embossed with a worker cell pattern and coated with a thin layer of beeswax (Fig. 1A-i,ii). Associated control electronics (including the detached scanner lid components) were housed in a separate, bee-free compartment at the top of the hive.

### 2.2 Artificial Comb Preparation

Given prior challenges with achieving consistent bee acceptance and comb adhesion directly on scanner surfaces, potentially due to material properties or heat sensitivity, the scanning interface for this experiment utilized indirectly mounted artificial cells. Attempts to cut and mount natural honeycomb cells onto a glass plate by melting their wax base (Fig. 1B-i) were also explored, but this process was too delicate, often resulting in structural damage to the cells. Subsequently, a 10×10 cm section, comprising of hexagonal cells (approx. 5.4 mm diameter) was 3D printed using Polymaker PolySmooth Transparent PVB filament (1.75 mm diameter). Following 3D printing, the comb section was subjected to three iterative cycles of immersion in isopropyl alcohol, with a drying period after each dip. This multi-step treatment was performed to smooth the surface by reducing the 3D-printed layer lines, a measure intended to minimize potential sites for bacterial settlement and thereby improve acceptance of the comb by the bees. The printed section was prepared for observation by first dipping it in diluted honey water to improve bee acceptance and allowing it to dry thoroughly. Subsequently, it was glued onto a separate 20×20 cm protective glass sheet (thickness: 0.8 mm) using a cyanoacrylate-based adhesive, as shown in Fig. 1B-ii. This sheet, carrying the 3D-printed cells, was positioned within the aperture in the wooden housing to sit directly on the scanner’s glass platen. Mounting the cell structure on the separate glass sheet prevented direct contact between the cells and the scanner surface. This arrangement also mitigated potential thermal effects from the scanner impacting the cell structure, its adhesion, or the bees inhabiting the cells, while providing a defined, bottomless cell structure for observation from beneath. A small access hole was maintained through the wooden housing structure above the scanner’s physical power button to permit manual restarts using a thin tool in the event of system crashes or power loss.

### 2.3 Control Software and Image Acquisition Setup

Scanner control on the SBC utilized the Linux SANE (Scanner Access Now Easy) interface [31], accessed via Epson’s proprietary “Epson Scan 2” software backend (v6.6.40.0 or later) [32] which provides necessary ARM-compatible drivers. Image acquisition was performed using the standard SANE utility “scanimage”. It was invoked by a custom Python 3 script which handled the precise scheduling of scans at regular intervals and managed the command-line parameters, including resolution and the specific ROI via cropping parameters. The direct use of the “scanimage” tool via the script was chosen for stability after initial tests with Python SANE library wrappers proved unreliable. Images were stored locally on the SBC’s SD card as files in a TIFF format and could be transferred via WiFi to network-attached storage. Manual restarts via the access hole were occasionally necessary following system interruptions.

### 2.4 Field Deployment and Data Acquisition Protocol

The BroodScan system was installed in an active honeybee colony (*A. mellifera*) housed at the apiary in the Botanical Garden of the University of Graz, Austria. To ensure targeted observation of brood development within the scanner’s field of view, the colony’s queen was confined to the 10×10 cm 3D-printed cell area (prepared as described in Section 2.2) using a standard queen excluder cage immediately after the scanner frame was inserted into the hive on 5 August 2022.

The initial period from 5 August to 9 August 2022 was primarily dedicated to system stabilization in the hive environment and iterative adjustments of the scanning parameters. While images were captured during this phase to monitor the condition of the comb and the queen’s initial activity (including the “Day 0 / August 5, 2022” state depicted in Figure 2A), settings for the Region of Interest (ROI), scan resolution, and output image format were still being finalized.

**Figure 2:**
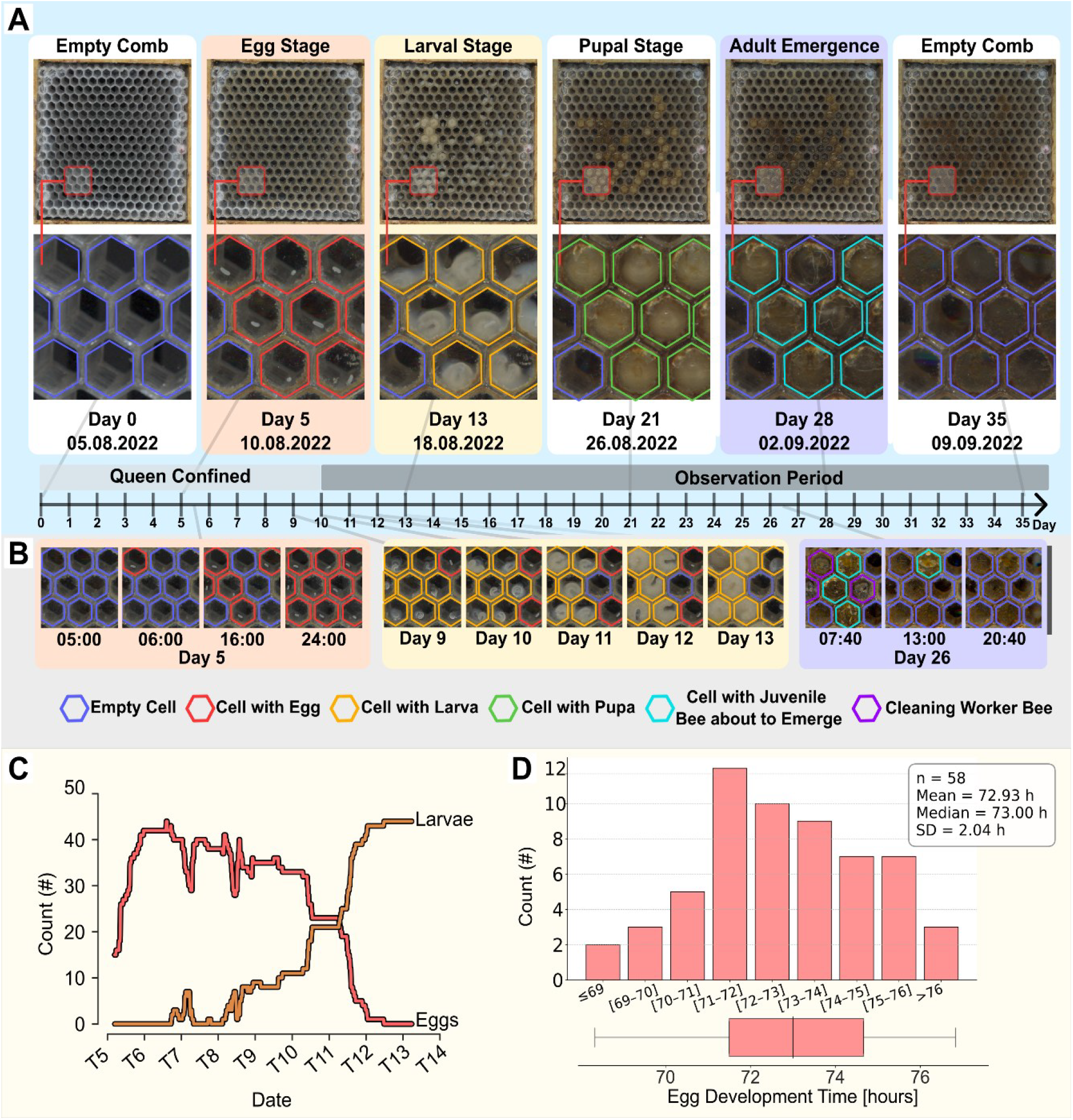
Observations over a complete cycle of honeybee brood development. Data were collected over 35 days, including an initial 10-day queen confinement period followed by a 25-day observation period. **(A)** Progression of brood development stages within the comb, showing global views of the Region of Interest and corresponding close-up sections at key time points (Days 0, 5, 13, 21, 28, 35). Stages progress from an empty comb, through egg, larval, and pupal stages, to adult emergence, eventually returning to an empty comb. **(B)** Close-up views highlighting key phases: egg deposition (e.g., on Day 5), larval growth (Days 9-13), and adult emergence (e.g., on Day 26). Cell contents are indicated by colored outlines (see legend). **(C)** Temporal dynamics of egg and larval counts over time (Days 5-14), derived exclusively from a cohort of cells that were successfully tracked through to adult bee emergence. This illustrates the egg-to-larva transition for brood that completed development. **(D)** Histogram showing the distribution of egg stage duration in hours for the cohort of 58 eggs that successfully developed through to adult emergence. The mean development time from oviposition to hatching was 72.93 ± 2.04 h (SD), with a median of 73.00 h.

Consistent, automated image acquisition with the optimized parameters detailed below commenced on 10 August 2022. For this main observation period (starting 10 August 2022), high-resolution scans were captured automatically every 20 minutes. The 10×10 cm ROI containing the cell structure was centered on the scanner platen to minimize parallax effects and scanned at 1200 dpi, resulting in 4992×4992 pixel images. Each scan took approximately 72 seconds to complete using the control software detailed in Section 2.3.

The queen remained caged on the observation area for approximately 10 days and was released back into the colony on 15 August 2022. Automated scanning continued, although the data collection experienced two significant interruptions (18–22 August and 2–8 September 2022) due to system halts requiring manual intervention. The overall observation period for this preliminary study concluded on 12 September 2022.

### 2.5 Image Annotation and Quantitative Data Extraction

A custom Python script was developed to facilitate frame-by-frame annotation of cell contents and key developmental events, utilizing the OpenCV library for core image manipulation and user interface elements. The imaging system allows for broad qualitative assessment of various cell states – such as identifying empty cells, or cells containing eggs, larvae, pupae, or adult bees engaged in activities like cell cleaning (as showcased in Figure 2A, B). To derive quantitative data from the acquired image sequences (for egg-to-larva development depicted in Figure 2C, D), a manual annotation process was employed. For this analysis, only cells with successfully emerging adult bees were considered. The presence of an egg within such a cell and its subsequent hatching and development into a larva over a period of several days was annotated across 1057 images.

These annotations were then processed to calculate: (1) the number of eggs, generating time-series counts; and (2) the egg development time.

### 2.6 *Varroa* Image Processing Pipeline

Due to the low contrast between *Varroa* mites and the maturing honeybee pupa, images used for visualizing mite development (Figure 3) required significant enhancement. A custom image processing pipeline was developed in Python (utilizing libraries such as OpenCV and NumPy) to improve mite visibility. This pipeline involved two parallel processing paths applied to the original color images:

**Figure 3:**
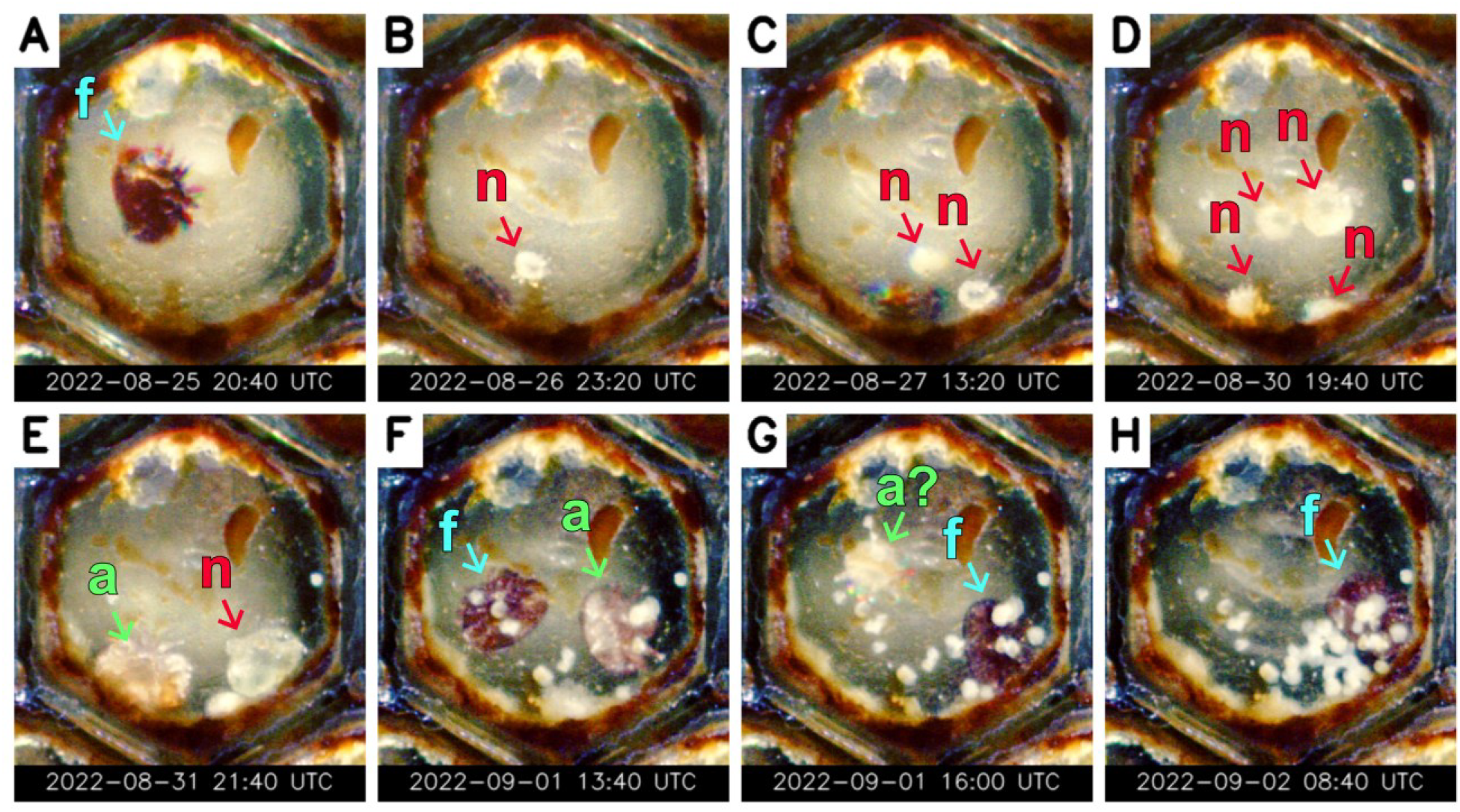
BroodScan imagery documenting Varroa destructor reproduction within a sealed worker cell. In the panels, colored arrows indicate representative mites: ‘f’ denotes the foundress mite (cyan arrow), ‘n’ denotes a nymph (red arrow), and ‘a’ denotes a newly molted/sclerotizing adult offspring (green arrow). Panels (A-H) show progression from (A) initial foundress sighting (Day ∼4 post-capping), (B-D) appearance and growth of offspring nymphs, (E) start of daughter sclerotization (first adult daughter), (F) near-mature daughter comparison with foundress, (G) plausible adult male presence, to (H) late-stage infestation prior to bee emergence (∼Day 12 post-capping). Note the persistent half-moon-shaped artifact in the upper right quadrant of the panels. Timestamps shown in panels are UTC.

1. Path A applied global contrast stretching, with limits determined by a specified saturation percentage of the overall image set’s intensity histogram.
2. Path B first applied per-channel (RGB) histogram equalization, followed by an increase in color saturation.

The images resulting from Path A and Path B were then blended with equal weighting. Finally, an unsharp masking filter was applied to the blended image to enhance fine details. Specific parameters for each step (e.g., saturation cut-off, saturation scaling factor, blending alpha, sharpening sigma and amount) were optimized empirically for best visual clarity.

## 3 RESULTS

Preliminary observational data were collected during the summer of 2022 using the BroodScan system and the experimental protocol detailed in Section 2. While operational interruptions during this initial field deployment (see Section 2.4) prevented the continuous tracking of any single individual bee through its entire development, the procedure successfully yielded high-resolution time-lapse imagery. This dataset captures examples of all honeybee developmental stages (egg, larva, pupa), key intra-cellular events (e.g., egg laying, hatching, larval growth, pupation, adult emergence, larval removal), and detailed observations of the intra-cellular reproductive cycle of the parasitic mite *Varroa destructor*. The following subsections present initial findings from these observations, focusing first on the overall honeybee brood development cycle (Section 3.1) and then on *Varroa* mite reproduction (Section 3.2).

### 3.1 Monitoring the Honeybee Brood Development Cycle

Using the BroodScan system, the complete honeybee brood development cycle, from egg laying to adult emergence, was monitored over a 35-day observation period, with queen confinement to the observation area for the initial 10 days to ensure oviposition. Figure 2 consolidates key observations and quantitative data derived from this monitoring, illustrating various developmental stages (subfigures A and B) and provides quantitative analyses of selected parameters, such as temporal changes in egg and larval counts (subfigure C) and the distribution of egg development times (subfigure D). Subfigure A provides a qualitative overview of the entire brood development cycle within the observation comb. Starting from an empty comb (Day 0), it shows the progression through queen oviposition (Egg Stage, Day 5), larval growth (Larval Stage, Day 13), the capped pupal stage (Pupal Stage, Day 21), and subsequent adult emergence (Adult Emergence, Day 28). By the end of the observation period (Day 35), the cells in the initially observed cohort had largely returned to an empty state following the completion of one brood cycle. The timeline inset in subfigure A indicates the queen confinement period and the overall observation period. Subfigure B highlights specific dynamic events within a selected close-up. It showcases rapid egg-laying by the queen on Day 5, where an initially empty area at 05:00 contained nine eggs by 24:00. The subfigure further illustrates larval growth and development between Day 9 and Day 13. Finally, on Day 26, the same close-up shows adult bees emerging from their cells.

Quantitative analysis of the annotated image sequences is presented in subfigure C, which displays the temporal dynamics of cell occupancy by tracking the number of cells containing eggs versus larvae within the close-up view from Day 5 to Day 14. The graph illustrates the initial increase in egg count, followed by a decline as eggs hatched, corresponding with a concurrent rise in the larval count. This demonstrates the clear transition wave from egg to larval stages within the observed cohort of cells. Subfigure D presents a histogram detailing the distribution of egg stage duration for a sample of 58 eggs. The development time, defined as the period from oviposition to larval hatching, had a mean of 72.93 ± 2.04 hours (Standard Deviation), with a median of 73.00 hours. The distribution of these egg development times appears approximately normal, centered around the 73-hour median.

### 3.2 Tracking Reproduction Intra-Cellular *Varroa*

The BroodScan system’s continuous, bottom-view monitoring allowed for detailed observation not only of honeybee brood development but also the typically hidden reproductive cycle of its parasite, *Varroa destructor*. We tracked this cycle within a single worker cell where the host bee egg (laid 13 Aug 2022, 13:10 UTC; hatched 16 Aug 14:10 UTC) was capped around 21 Aug PM. Developmental timings hereafter are given in **Post-Capping Days (PCD)** relative to this capping estimate. Because visualizing the mites against the maturing honeybee pupa (which forms the cell background in our images) can be challenging, the images in Figure 3 were enhanced using a blend of contrast stretching and per-channel histogram equalization (see Section 2.6 for details). Note the persistent presence of a small, half-moon-shaped artifact in the upper right quadrant throughout the sequence (visible in Fig. 3A-H); its appearance suggests it might be a piece of propolis or other hive debris lodged in the cell near the time of capping. The key developmental stages observed were:

1. **Foundress Mite Visible (∼PCD 4):** The dark reddish-brown foundress mite is clearly visible at the cell base by 25 Aug (Fig. 3A). Having necessarily entered the cell before it was capped (∼PCD 0), this marks the first point she was clearly distinguishable in the recordings. The background host prepupa/early pupa is pale.
2. **First Offspring (∼PCD 5):** A small, pale nymph (presumed male, based on laying sequence), appears alongside the foundress (Fig. 3B). This timing aligns with the male egg hatching.
3. **Second Offspring (∼PCD 6):** A second pale nymph (presumed first daughter) is distinguishable (Fig. 3C), consistent with the expected hatching time of the second egg laid (∼30 hours after the first).
4. **Multiple Offspring Visible (∼PCD 9):** Population growth is evident by 30 Aug, with at least three distinct pale offspring mites seen simultaneously (Fig. 3D). Guanine deposits (white spots) become noticeable.
5. **Daughter Sclerotization Begins (∼PCD 10):** Around 31 Aug (Fig. 3E), a larger offspring (presumed first daughter) begins showing light brown coloration, indicating the start of post-molt sclerotization into a young adult female. Another large pale mite is present. The host pupa background starts showing slight darkening.
6. **Near-Mature Daughter (∼PCD 11):** By 01 Sep (Fig. 3F), the sclerotizing daughter has darkened significantly, appearing medium/dark brown, similar to the foundress (also visible). Guanine is prominent. The host pupa background now appears distinctly darker/browner.
7. **Potential Adult Male Sighted (∼PCD 11):** Shortly later, alongside the foundress, a distinct small, pale mite is visible (Fig. 3G), plausibly the mature male required for mating.
8. **Final Observation (∼PCD 12):** The last recorded image on 02 Sep (08:40 UTC), taken before monitoring ceased, clearly shows the foundress mite and one other smaller, pale mite (nymph or potential male) amidst heavy guanine accumulation (Fig. 3H). The host pupa background is dark brown, indicating imminent bee emergence.

This sequence (Fig. 3), captured via the BroodScan’s unique basal perspective, provides a rare, detailed visual record of *Varroa destructor*’s reproductive cycle unfolding within a sealed cell. We were able to track the sequential appearance of offspring, observe the distinct process of daughter mite sclerotization and darkening, and monitor the accumulation of guanine deposits. Concurrently, the visible darkening of the host pupa served as an internal developmental clock for the observed mite milestones. These observations vividly document processes typically obscured beneath the wax capping.

The continuous monitoring capability was crucial for capturing these dynamic events. Unfortunately, a system interruption on PCD 12 (02 Sep) – the presumed day of host emergence – prematurely terminated data collection for this cell. This prevented direct observation of crucial final events: the bee’s emergence, the number of mites departing with the host, and the fate of any mites remaining within the cell. Nevertheless, the captured data leading up to this point offers valuable insights into the timing and visual markers of intra-cellular *Varroa* reproduction.

## 4 DISCUSSION

Investigations into the intra-cellular dynamics of honeybee brood development, especially within sealed cells for studies on parasites like *Varroa destructor*, other diseases, or the normative development of honeybee brood itself, have historically been constrained by destructive methodologies. These typically involve the manual uncapping and excavation of individual cells, a process that sacrifices the developing bee and inherently prohibits continuous, non-invasive observation [12–13, 16–17, 33]. The BroodScan technology, presented in this paper, details experiments conducted in 2022 and builds upon core concepts first publicly documented with the Hiveopolis project since 2022 [23–27]. Our study investigates this observational avenue, critically employing a CCD-based system specifically for high-resolution basal monitoring. Our preliminary findings successfully demonstrate BroodScan’s capacity to observe the complete honeybee brood development cycle from egg to adult emergence (Section 3.1, Figure 2) and, notably, to provide detailed visual documentation of the intra-cellular reproductive cycle of the parasitic mite *Varroa destructor* within these sealed cells (Section 3.2, Figure 3). The recent work by Borlinghaus et al. (2024) [34], employing a CIS-based flatbed scanner for in-hive monitoring of brood and pathogens during their 2024 field season, has affirmed the potential of this general scanning approach to overcome some of these limitations.

The BroodScan system was specifically designed to overcome several critical limitations inherent in traditional top-down observation methods. These traditional approaches are often hampered by disruptive frame removal and visual obstructions from wax cappings and bee traffic (Section 1.2, Figure 1C). By imaging through a transparent glass base upon which bottomless 3D-printed cells are placed, BroodScan provides an unobstructed, close-distance, high-resolution view of intra-cellular contents. A key technological distinction of our BroodScan system is the use of a CCD-based flatbed scanner, chosen deliberately over CIS alternatives due to the CCD’s superior depth of field. This characteristic is essential for resolving three-dimensional details within the honeycomb cells, particularly for observing developing pupae and mites situated at varying depths within the cell (as detailed in Section 2.1). While flatbed scanner technology, in general, offers advantages in reducing geometric distortions compared to some camera-based systems viewing wide areas at close proximity, the selected CCD scanner does exhibit some parallax, especially towards its lateral edges. This was mitigated by centering the 10×10 cm Region of Interest (ROI) (Section 2.1). This strategy represents a conscious trade-off: prioritizing the crucial depth of field required for detailed in-cell imaging with CCD technology over the near-zero parallax but more limited depth of field typically associated with CIS scanners.

Our observations of the honeybee brood development cycle yielded quantitative data, such as an average egg development time of approximately 73±2 hours (Figure 2D), obtained non-destructively. The ability to continuously monitor individual cells allows for the precise timing of key developmental events like hatching and pupation, and the tracking of cell content changes over time (Figure 2C). This contrasts sharply with methods requiring repeated cell uncapping or comb destruction, which can alter the natural progression of development and are labor-intensive [12–13, 16–17, 33]. While other automated hive monitoring systems exist, they typically focus on colony-level parameters (e.g., weight, temperature, acoustics) or tracking adult bee activity at the entrance or on the comb surface using cameras or RFID [35–38]. Repurposed flatbed scanner technology has been utilized for biological time-lapse imaging in diverse fields, for example, to monitor nematode populations [39], bacterial swarming [40], automated monitoring of insect larvae [41] and plant root dynamics [42]. In contrast, the BroodScan system represents a specific adaptation of this technology, developed for the detailed, longitudinal, and *in situ* investigation of intra-cellular processes as they occur within the environment of fully operational honeybee hives.

One of the most compelling demonstrations of BroodScan’s utility is its application to visualizing *Varroa destructor* reproduction within sealed worker brood cells (Figure 3). This parasite represents a major threat to honeybee health [MS-B05], and much of its critical reproductive phase occurs hidden beneath the wax capping, making it notoriously difficult to study *in situ* without invasive techniques [12–13]. We successfully monitored *Varroa* presence and offspring (primarily in drone brood) through the CCD-based BroodScan system, leveraging its superior depth of field and a dedicated image enhancement pipeline (Section 2.6), facilitated a particularly granular documentation of the mite’s reproductive stages within worker cells. Specifically, we were able to clearly observe the initial foundress mite, the sequential appearance and development of distinct nymphal stages, the characteristic sclerotization of daughter mites as they matured, and the accumulation of guanine deposits, all while correlating these mite-specific events with the visible maturation of the host pupa (as depicted in Figure 3). Such detailed visual timelines of the complete mite reproductive cycle offer considerable potential for understanding host– parasite interactions at an intra-cellular level, for precisely timing mite developmental phases, and for more direct assessment of the efficacy of *Varroa* control measures that target these cryptic stages.

The BroodScan system leverages consumer-grade scanner technology, importantly, its relatively low cost, aligning with a broader trend of ‘frugal science’ approaches that adapt accessible tools for complex biological research [15, 41, 43]. This cost-effectiveness enhances its potential for wider adoption. Further strengths include its potential for autonomous long-term data acquisition with minimal disturbance once installed, and the high spatial resolution and good geometric fidelity it offers for events occurring at the base of the cells. The direct view from beneath circumvents the significant observational hurdle of cell cappings, opening up the entire pupal stage, and *Varroa*’s reproductive phase, to continuous visual scrutiny.

Despite these promising initial results, the current BroodScan prototype and deployment have several limitations that warrant discussion, some of which may be inherent to adapting current consumer-grade scanner technology for complex in-hive monitoring. Firstly, the 10×10 cm observation area, while providing a significant number of cells for study, represents only a fraction of a standard brood frame. Although full-frame 3D-printed comb sections are technically feasible to fabricate and are generally well accepted by bees, significantly expanding the *active scanning area* poses distinct challenges for the current BroodScan system, particularly given its reliance on a consumer-grade CCD scanner. A larger scanned area would proportionally increase the duration of disturbance to the colony during each scan (i.e., prolonged movement of the bright CCD scan bar, associated noise, and vibrations), potentially altering bee behavior or the brood nest environment and thus counteracting the goal of minimally invasive observation. Specifically, horizontal expansion of the scanned ROI would exacerbate the existing parallax distortions observed towards the scanner’s lateral edges (Section 2.1). Both horizontal expansion and attempts to further increase resolution would also lead to considerably longer scan times (currently ∼70 seconds for the 10×10 cm ROI at 1200 dpi, yielding ∼70 MB TIFF files) and correspondingly larger data volumes, posing practical limits for data handling and storage in extended deployments.

Secondly, specific aspects of the materials used and operational factors in this preliminary study require further consideration for long-term deployments. While the 3D-printed PVB cells, treated with honey water, demonstrated good acceptance by the bees during the observation period of this study (see Section 2.2), comprehensive research into the long-term effects of this specific material on cell microclimate, its durability against repeated cleaning cycles or hive debris accumulation, and any potential impact on bee physiology or behavior over extended periods is still required. The field deployment also experienced operational interruptions (Section 2.4), highlighting the need for further refinement to ensure robust, continuous operation for multi-season studies. While the CCD scanner offered good depth of field, some parallax effects were still noted towards the edges of the platen (Section 2.1), necessitating careful ROI selection. The current extraction of quantitative data also relies on manual annotation (Section 2.5), which is labor-intensive and may limit the scale of analysis. Automation of image analysis for recognizing cell contents and events is therefore a critical next step. Finally, while white light illumination used in this study showed no obvious detrimental effects on brood development (Section 1.4), future iterations could incorporate red-light illumination, to which bees are largely insensitive, to further minimize any potential disturbance.

Future development of the BroodScan technology could focus on addressing the limitations identified, thereby expanding its capabilities and research applications. To overcome the current constraints on observation area, approaches such as exploring larger format scanners or, inspired by systems like the *C. elegans* Lifespan Machine [39], tiling multiple synchronized scanning units could be investigated. Enhancing overall system robustness for multi-season, autonomous deployment remains a key objective. Crucially, the development of automated image analysis algorithms, potentially employing machine learning techniques [34, 43], is essential for efficiently processing the large datasets generated and for extracting quantitative data on a larger scale than currently possible with manual annotation. The systems utility extends beyond fundamental studies in honeybee developmental biology (Section 3.1) and host–parasite interactions with *Varroa destructor* (Section 3.2). The capacity for continuous, in-situ monitoring with the BroodScan system enables detailed investigation into the sub-lethal effects of environmental stressors, such as pesticides, varied nutritional inputs, or temperature fluctuations on brood health, developmental trajectories, and associated bee behaviors in real-time. Such an approach is poised to yield more direct and nuanced insights than studies relying primarily on colony-level outcomes or destructive sampling techniques [39–40].

The performance and design compactness of systems like BroodScan stand to benefit considerably from ongoing advancements in sensor technology. To illustrate, current Contact Image Sensor (CIS) technology, despite advantages in low parallax and size, struggles with the limited depth of field necessary for imaging 3D honeycomb structures. Nevertheless, future CIS or other sensor technologies that achieve improved depth of field while maintaining these desirable characteristics could pave the way for even more streamlined and energy-efficient brood scanning solutions.

In conclusion, the BroodScan system, by innovatively repurposing accessible flatbed scanner technology as demonstrated in these preliminary experiments, offers a novel and valuable method for obtaining high-resolution, longitudinal data on previously obscured intra-cellular processes within the honeybee brood nest. It successfully overcomes significant limitations of traditional observational techniques and opens crucial new avenues for research into honeybee developmental biology, colony health, intricate host–parasite interactions like those involving the *Varroa* mite, and the impacts of various environmental stressors. As technologies like BroodScan mature, offering a continuous window into the hive at the cellular level, they will become increasingly instrumental not only for fundamental apicultural research but also for the practical stewardship and conservation of honeybee populations in changing environments.

## ACKNOWLEDGEMENTS

We thank Pollenity for their expert support in 3D printing and for supplying the 3D-printed combs. We are also grateful to Sarah Schönwetter-Fuchs for the essential groundwork that underpinned the investigations reported in this paper.

## REFERENCES

[1] Zion Market Research. Honey Market By Region - Global And Regional Industry Overview, Market Intelligence, Comprehensive Analysis, Historical Data, And Forecasts 2024 – 2032. Zion Market Research (2024). Available at: https://www.zionmarketresearch.com/report/honey-market (Accessed: 15 May 2025).

[2] Gallai, N., Salles, J. M., Settele, J. & Vaissière, B. E. Economic valuation of the vulnerability of world agriculture confronted with pollinator decline. Ecological Economics 68, 810–821. doi:10.1016/j.ecolecon.2008.06.014 (2009).

[3] Klein, A. M., Vaissière, B. E., Cane, J. H., Steffan-Dewenter, I., Cunningham, S. A., Kremen, C. & Tscharntke, T. Importance of pollinators in changing landscapes for world crops. Proceedings of the Royal Society B: Biological Sciences 274, 303–313. doi:10.1098/rspb.2006.3721 (2007).

[4] VanEngelsdorp, D., Hayes Jr, J., Underwood, R. M. & Pettis, J. A survey of honey bee colony losses in the US, fall 2007 to spring 2008. PLOS ONE 3(12), 1–6. doi:10.1371/journal.pone.0004071 (2008).

[5] Neumann, P. & Carreck, N. L. Honey bee colony losses. Journal of Apicultural Research 49(1), 1–6. doi:10.3896/IBRA.1.49.1.01 (2010).

[6] Seitz, N., Traynor, K. S., Steinhauer, N., Rennich, K., Wilson, M. E., Ellis, J. D., Rose, R., Tarpy, D. R., Sagili, R. R., Caron, D. M., Delaplane, K. S., Rangel, J., Lee, K., Baylis, K., Wilkes, J. T., Skinner, J. A., Pettis, J. S. & vanEngelsdorp, D. A national survey of managed honey bee 2014–2015 annual colony losses in the USA. Journal of Apicultural Research 54(4), 292– 304. doi:10.1080/00218839.2016.1153294 (2022).

[7] Gray, A., Adjlane, N., Arab, A., Ballis, A., Brusbardis, V., Bugeja Douglas, A., Cadahía, L., Charrière, J.-D., Chlebo, R., Coffey, M. F., Cornelissen, B., Amaro da Costa, C., Danneels, E., Danihlík, J., Dobrescu, C., Evans, G., Fedoriak, M., Forsythe, I., Gregorc, A., Arakelyan, I. I., Johannesen, J., Kauko, L., Kristiansen, P., Martikkala, M., Martín-Hernández, R., Mazur, E., Medina-Flores, C. A., Mutinelli, F., Omar, E. M., Patalano, S., Raudmets, A., San Martin, G., Soroker, V., Stahlmann-Brown, P., Stevanovic, J., Uzunov, A., Vejsnaes, F., Williams, A. & Brodschneider, R. Honey bee colony loss rates in 37 countries using the COLOSS survey for winter 2019–2020: the combined effects of operation size, migration and queen replacement. Journal of Apicultural Research 62(2), 204–210. doi:10.1080/00218839.2022.2113329 (2023).

[8] Nazzi, F. & Le Conte, Y. Ecology of Varroa destructor, the major ectoparasite of the western honey bee, Apis mellifera. Annual Review of Entomology 61(1), 417–432. doi:10.1146/annurev-ento-010715-023731 (2016).

[9] Stephan, J. G., de Miranda, J. R. & Forsgren, E. American foulbrood in a honeybee colony: spore-symptom relationship and feedbacks between disease and colony development. BMC Ecology 20, 15. doi:10.1186/s12898-020-00283-w (2020).

[10] Heath, L. A. F. Development of chalk brood in a honeybee colony: a review. Bee World 63(3), 119–130. doi:10.1080/0005772X.1982.11097876 (1982).

[11] Winston, M. L. The Biology of the Honey Bee. (Harvard University Press, 1991).

[12] Dietemann, V., Nazzi, F., Martin, S. J., Anderson, D. L., Locke, B., Delaplane, K. S., Wauquiez, Q., Tannahill, C., Frey, E., Ziegelmann, B., Rosenkranz, P. & Ellis, J. D. Standard methods for varroa research. Journal of Apicultural Research 52(1), 1–54. doi:10.3896/IBRA.1.52.1.09 (2013).

[13] Uzunov, A., Janashia, I., Chen, C., Costa, C. & Kovačić, M. A scientific note on ‘Rapid brood decapping’—a method for assessment of honey bee (Apis mellifera) brood infestation with Tropilaelaps mercedesae. Apidologie 56(2), 40. doi:10.1007/s13592-025-01171-2 (2025).

[14] Erickson, E. H. Stress and honey bees. Gleanings in Bee Culture 118(11), 650–654. (1990).

[15] Göröcs, Z. & Ozcan, A. Biomedical imaging and sensing using flatbed scanners. Lab on a Chip 14(17), 3248–3257. doi:10.1039/C4LC00530A (2014).

[16] Human, H., Brodschneider, R., Dietemann, V., Dively, G., Ellis, J. D., Forsgren, E., Fries, I., Hatjina, F., Hu, F.-L., Jaffé, R., Jensen, A. B., Köhler, A., Magyar, J. P., Özkýrým, A., Pirk, C. W. W., Rose, R., Strauss, U., Tanner, G., Tarpy, D.R., van der Steen, J. J. M., Vaudo, A., Vejsnæs, F., Wilde, J., Williams, G. R. & Zheng, H. Q. Miscellaneous standard methods for Apis mellifera research. Journal of Apicultural Research 52(4), 1– 53. doi:10.3896/IBRA.1.52.4.10 (2013).

[17] Oertel, E. Metamorphosis in the honeybee. Journal of Morphology 50(2), 295–339. doi:10.1002/jmor.1050500202 (1930).

[18] Stabentheiner, A., Kovac, H. & Brodschneider, R. Honeybee colony thermoregulation – regulatory mechanisms and contribution of individuals in dependence on age, location and thermal stress. PLOS ONE 5(1), e8967. doi:10.1371/journal.pone.0008967 (2010).

[19] Schultner, E., Oettler, J. & Helanterä, H. The role of brood in eusocial Hymenoptera. The Quarterly Review of Biology 92(1), 39–78. doi:10.1086/690840 (2017).

[20] Webb, J. N., Houston, A. I., McNamara, J. M. & SzéKely, T. Multiple patterns of parental care. Animal Behaviour 58(5), 983–993. doi:10.1006/anbe.1999.1215 (1999).

[21] Cridge, A. G., Lovegrove, M. R., Skelly, J. G., Taylor, S. E., Petersen, G. E. L., Cameron, R. C. & Dearden, P. K. The honeybee as a model insect for developmental genetics. genesis 55(5), e23019. doi:10.1002/dvg.23019 (2017).

[22] Naiem, E. S., Hrassnigg, N. & Crailsheim, K. Nurse bees support the physiological development of young bees (Apis mellifera L.). Journal of Comparative Physiology B 169, 271–279. doi:10.1007/s003600050221 (1999).

[23] HIVEOPOLIS. Final prototype of the brood nest module with integrated camera system. Tech. Rep. D5.3 (The European Union, 2022). doi:10.3030/824069 (confidential)

[24] HIVEOPOLIS. Report on the performance of HIVEOPOLIS bio-hybrids. Tech. Rep. D2.6 (The European Union, 2024). doi:10.3030/824069

[25] HIVEOPOLIS. Prototype of the complete system. Tech. Rep. D2.5 (The European Union, 2023). doi:10.3030/824069

[26] HIVEOPOLIS. Report on automated brood assessment & affection system. Tech. Rep. D5.4 (The European Union, 2023). doi:10.3030/824069 (confidential)

[27] HIVEOPOLIS. Final report on community forming. Tech. Rep. D7.4 (The European Union, 2024). doi:10.3030/824069

[28] TWAIN Working Group. TWAIN Standard. Available: https://twain.org/. Accessed: May 9, 2025.

[29] Lichtenstein, L., Grübel, K. & Spaethe, J. Opsin expression patterns coincide with photoreceptor development during pupal development in the honey bee, Apis mellifera. BMC Developmental Biology 18, 1. doi:10.1186/s12861-018-0162-8 (2018).

[30] Burke, B., Jorden, P. & Vu, P. CCD Technology. Experimental Astronomy 19, 69–102. doi:10.1007/s10686-005-9011-4 (2005).

[31] SANE Project. Scanner Access Now Easy. Available: http://www.sane-project.org/. Accessed: May 9, 2025.

[32] Epson Corporation. Epson Scan 2 Software and Drivers (Version 6.6.40.0 or later). Available: https://support.epson.net/linux/en/epsonscan2.php. Accessed: May 9, 2025.

[33] Fleig, R. & Sander, K. Honeybee morphogenesis: embryonic cell movements that shape the larval body. Development 103(3), 525– 534. doi:10.1242/dev.103.3.525 (1988).

[34] Borlinghaus, P., Gülzow, J. M. & Odemer, R. In-hive flatbed scanners for non-destructive, long-term monitoring of honey bee brood, pathogens and pests. Smart Agricultural Technology 9, 100655. doi:10.1016/j.atech.2024.100655 (2024).

[35] Ulrich, J., Stefanec, M., Rekabi-Bana, F., Fedotoff, L. A., Rouček, T., Gündeğer, B. Y., Saadat, M., Blaha, J., Janota, J., Hofstadler, D. N., Žampachů, K., Keyvan, E. E., Erdem, B., Şahin, E., Alemdar, H., Turgut, A. E., Arvin F., Schmickl, T. & Krajník, T. Autonomous tracking of honey bee behaviors over long-term periods with cooperating robots. Science Robotics 9(95), eadn6848. doi:10.1126/scirobotics.adn6848 (2024).

[36] Ai, H. & Takahashi, S. The lifelog monitoring system for honeybees: RFID and camera recordings in an observation hive. Journal of Robotics and Mechatronics 33(3), 457–465. doi:10.20965/jrm.2021.p0457 (2021).

[37] Meikle, W. G., Holst, N., Colin, T., Weiss, M., Carroll, M. J., McFrederick, Q. S. & Barron, A. B. Using within-day hive weight changes to measure environmental effects on honey bee colonies. PLOS ONE 13(5), e0197589. doi:10.1371/journal.pone.0197589 (2018).

[38] Bozek, K., Hebert, L., Portugal, Y., Mikheyev, A. S. & Stephens, G. J. Markerless tracking of an entire honey bee colony. Nature Communications 12(1), 1733. doi:10.1038/s41467-021-21769-1 (2021).

[39] Stroustrup, N., Ulmschneider, B. E., Nash, Z. M., López-Moyado, I. F., Apfeld, J. & Fontana, W. The Caenorhabditis elegans lifespan machine. Nature Methods 10, 665–670. doi:10.1038/nmeth.2475 (2013).

[40] Bru, J.-L., Siryaporn, A. & Høyland-Kroghsbo, N. M. Time-lapse imaging of bacterial swarms and the collective stress response. Journal of Visualized Experiments (JoVE) 159, e60915. doi:10.3791/60915 (2020).

[41] Ohki, H., Seong, K. H., Suzuki, T. & Shimoda, M. Automated survival monitoring system: exploring starvation resistance in newly hatched larvae. Applied Entomology and Zoology 60, 99– 108. doi:10.1007/s13355-025-00896-x (2025).

[42] Smith, H. C., Niewohner, D. J., Dewey, G. D., Longo, A. M., Guy, T. L., Higgins, B. R., Daehling, S. B., Genrich, S. C, Wentworth, C. D. & Durham Brooks, T. L. Using flatbed scanners to collect high-resolution time-lapsed images of the Arabidopsis root gravitropic response. Journal of Visualized Experiments (JoVE) 83, e50878. doi:10.3791/50878 (2014).

[43] Ong, S.-Q., Pinoy, N., Lim, M. H., Bjerge, K., Peris-Felipo, F. J., Lind, R., Cuff, J. P., Cook, S. M. & Høye, T. T. ScannerVision: Scanner-based image acquisition of medically important arthropods for the development of computer vision and deep learning models. Current Research in Parasitology & Vector-Borne Diseases 7, 100268. doi: 10.1016/j.crpvbd.2025.100268 (2025).

